# Repeated heat waves trigger divergent transcriptional responses in cold- and warm-adapted yeast species

**DOI:** 10.64898/2026.02.02.703184

**Authors:** Chloé Haberkorn, Jennifer Molinet, Rike Stelkens

**Affiliations:** Department of Zoology, Stockholm University, Svante Arrheniusväg 18 B, SE-10691 Stockholm, Sweden; Facultad de Ingeniería, Instituto de Ciencias Aplicadas, Centro de Investigación e Innovación, Universidad Autónoma de Chile, Santiago, Chile; ANID-Millennium Science Initiative-Millennium Institute for Integrative Biology (iBio), Santiago, Chile

**Keywords:** heat wave, yeast, thermal tolerance, transcriptional memory

## Abstract

Extreme climatic events such as heat waves pose major challenges to species survival and have profound impacts on evolutionary processes. Plasticity is thought to buffer organismal stress, yet the molecular mechanisms underlying repeated stress responses, especially the roles of transcriptional plasticity and stress memory, remain mostly unresolved. Here, we experimentally exposed clonal populations of eight divergent *Saccharomyces* yeast species with different thermal tolerances to repeated heat waves and compared their phenotypic and transcriptomic profiles after a few generations of mitotic growth. Warm-adapted species maintained higher growth than cold-adapted species across heat wave exposures. Thermo-generalist species showed intermediate outcomes with one species improving growth across repeated heat waves. Using a conceptual framework separating no-memory (gene expression independent from prior exposure) from memory-like responses (expression modulated by prior exposure) showed mostly no-memory responses with conserved transcriptomic signatures of proteostasis induction, and reduced expression of ribosome biogenesis and translation upon repeated heat waves. Memory-like responses were highly species-specific, revealing strikingly opposite patterns of (de)sensitization in ribosomal and translational pathways in species at the two extremes of thermal tolerance. Together, our results show that thermal resilience in yeast is associated with alternative transcriptional changes and suggest that warm-and cold-temperature specialists use divergent gene regulatory strategies upon repeated heat waves. With climate change projected to increase the frequency and intensity of heat waves, understanding plastic responses across species with diverse ecological and genetic backgrounds is crucial for predicting biological resilience.

## Introduction

Extreme climatic events, such as heat waves, are increasingly recognized as major drivers of ecological and evolutionary change. Heat waves can be defined as “events in which temperatures are excessively higher than normal for several consecutive days” (Domeisen *et al*., 2023), i.e., a repeated exposure to heat shock. Current climate simulations project that heat waves will become more frequent, with a concomitant rise in intensity and duration (Domeisen *et al*., 2023). Heat waves impose rapid and intense challenges to cellular function, physiology, and survival, frequently pushing organisms close to or beyond their thermal tolerance limits, which can have lethal or sublethal effects on natural populations (Hemraj *et al*., 2020; Breshears *et al*., 2021). For instance, simulated heat wave exposure in the three-spined stickleback reduced reproductive success (Barrett and Stein, 2024) and offspring survival (Spence-Jones *et al*., 2025) and induced long-lasting immune impairments (Dittmar *et al*., 2014), increasing susceptibility to infectious disease and facilitating pathogen spread. Bird species with small thermal ranges showed sharp declines in population growth rates after a heat wave (Jiguet *et al*., 2006). In anuran amphibians, heat waves were shown to affect larval stages by delaying metamorphosis and impacting sexual development, disrupting populational sex ratios (Ujszegi *et al*., 2022). In insects, heat waves have been shown to reduce fertility, alter reproductive traits, and impose demographic bottlenecks (Sales *et al*., 2018; Haque, Paul and Khan, 2025; Taig *et al*., 2025), even in the absence of lethal temperatures (Cao *et al*., 2025). At the microbial level, different studies highlight that climate change, notably through the increased frequency of heat waves, drought, and flooding, is driving the geographical expansion and thermotolerance of non-native fungal species (Porter, 1980; Deng *et al*., 2015; Friedman and Schwartz, 2019). This potentially alters ecosystem dynamics (Tischner *et al*., 2022) but also raises public health concerns, as previously harmless or geographically constrained fungi become opportunistic pathogens, for instance those involved in cutaneous (Gadre *et al*., 2022) or respiratory infections (Sondermeyer *et al*., 2016). In summary, heat wave effects have been documented across a broad range of taxa, revealing striking differences in vulnerability and resilience. However, cross-species comparisons remain rare, despite being essential for understanding how biodiversity will respond to increasingly frequent and intense climate extremes. A key unresolved question in the context of climate change is: Do species with different thermal tolerance profiles rely on common or distinct responses to cope with repeated heat waves, and can these responses be predicted from their thermal profiles? Such comparisons are challenging, as they typically require long-term monitoring of natural populations and comprehensive datasets that capture both organismal traits (e.g. growth and survival) and environmental variability.

Laboratory microbial model organisms provide powerful systems to investigate the molecular and transcriptomic basis of thermotolerance, as they enable controlled experimental comparisons across species. The genus *Saccharomyces* is well suited for such comparative approaches, as thermal tolerance has been extensively characterized across species (Gonçalves *et al*., 2011; Salvadó *et al*., 2011; Weiss *et al*., 2018; Abrams *et al*., 2021) revealing substantial interspecific variation and evolvability of thermal performance traits (Molinet and Stelkens, 2025). In addition, previous work has shown that transcriptional and metabolic responses to temperature are highly variable among strains and species (Caspeta Luis and Nielsen Jens, 2015; Fay *et al*., 2023). Together with the availability of high-quality genomes and annotations for all species (Bendixsen, Frazão and Stelkens, 2022), *Saccharomyces* yeast is an ideal system to investigate the molecular and phenotypic responses to heat waves, comparatively across a whole genus. Because of their short generation time (ca. 2.5h), temporal data can also easily be collected across multiple generations.

Responses to acute heat shock are relatively conserved across yeast species (Mühlhofer *et al*., 2019; Jann *et al*., 2020), triggering a rapid and coordinated induction of heat shock proteins (HSPs), which are central components of the cellular stress response (Mühlhofer *et al*., 2019). Heat shock can disrupt the balance between protein synthesis and proper folding, leading to the accumulation of misfolded proteins in the cell. HSPs, such as molecular chaperones, facilitate correct protein folding, thus buffering proteotoxic stress (Mühlhofer *et al*., 2019). While heat-shock responses in yeasts have been extensively characterised, the transcriptional dynamics associated with heat waves - repeated and prolonged heat events, representing an ecologically relevant climate scenario - have not been investigated. In particular, it is unclear whether heat waves engage distinct or additional gene regulatory networks beyond those involved in the heat-shock response. Repeated heat stress may induce acclimation, defined here as a change in the magnitude or direction of the response following previous exposure. In unicellular organisms, such responses may persist over several mitotic divisions and therefore resemble forms of transcriptional memory (Brickner *et al*., 2007; D’Urso and Brickner, 2014; D’Urso *et al*., 2016). Transcriptional memory has been described as the ability of prior stress exposure to alter the speed, magnitude, or persistence of gene expression upon subsequent stress encounters (Lämke and Bäurle, 2017; Bäurle and Trindade, 2020; Vihervaara *et al*., 2021). Such memory-like responses can arise from incomplete resetting of regulatory states, persistent chromatin marks, transcriptional bookmarking, or other mitotically inherited molecular states (Light *et al*., 2010; D’Urso and Brickner, 2014; Fitz-James and Cavalli, 2022; Pratx *et al*., 2023). However, whether repeated heat waves elicit conserved or species-specific transcriptional memory across closely related species remains largely unknown.

Here, we measured immediate transcriptional responses to repeated heat waves by analysing differential gene expression patterns across genetically and ecologically different species of *Saccharomyces* yeasts. We used large, diploid populations of 16 strains from eight species with vastly different thermal tolerances (*S. cerevisiae, S. paradoxus, S. mikatae, S. eubayanus, S. kudriavzevii, S. uvarum, S. arboricola*, and S. *jurei*) and exposed them for two hours to 37°C heat events, for five consecutive days, propagating populations with serial transfer. We selected 37°C as a standardized extreme condition to mimic heat wave exposure across species. This temperature elicits gene expression changes in all species, without compromising survival. We compared the transcriptomes of all populations before and after the first heat wave on day 1, and before and after repeated exposure to the heat wave on day 5. An 18-hour recovery period at 25°C between heat waves allowed populations to undergo approximately 17 generations of mitotic growth over the five days. Because the fixation of beneficial mutations and/or karyotypic changes usually require hundreds of generations, short-term changes in phenotypic performance occurring in less than 20 generations are expected to be driven predominantly by mitotically inherited transgenerational plasticity (D’Urso and Brickner, 2014). Through the analysis of differentially expressed genes, we assessed the cumulative effects of repeated heat waves over time, by disentangling (i) no-memory transcriptional responses, where gene expression was independent of prior exposure and (ii) memory-like responses, where gene expression was modulated (likely through epigenetic mechanisms) by prior exposure. We identified both conserved and divergent pathways between species with different thermal tolerances. Heat wave exposure consistently induced proteostasis pathways and repressed translation and ribosome biogenesis, mirroring the core features of the yeast heat-shock response. However, we also found highly species-specific memory-like transcriptional changes, indicating that species with different thermal tolerances modulate specific regulatory pathways upon repeated heat wave exposure.

## Material & Methods

### Saccharomyces strains

We used 16 strains, two from each of eight *Saccharomyces* species: *S. cerevisiae, S. paradoxus, S. mikatae, S. eubayanus, S. kudriavzevii, S. uvarum, S. arboricola*, and *S. jurei* (Table 1). Divergence times between these species are estimated to be between 2 million and 20 million years (Kellis *et al*., 2003; Borneman and Pretorius, 2015; Shen *et al*., 2018). *Saccharomyces* species vary widely in their temperature preferences, with some cryotolerant strains growing at 0°C while some thermotolerant strains can grow in temperatures as high as 42°C (Salvadó *et al*., 2011). Strains within species were isolated from different ecological niches (details in Supplementary Table S1). They differ genetically and phenotypically from each other (Liti *et al*., 2009; Warringer *et al*., 2011; Peter *et al*., 2018) to different extents, depending on the population structure and genetic diversity present within each species (ranging from only 0.4% sequence divergence within *S. cerevisiae* to ca 2% within *S. paradoxus*) (Peris *et al*., 2023). Here, we used the same 16 strains for which thermal performance curves have been accurately described recently (Molinet and Stelkens, 2025). *S. cerevisiae* is the most heat-tolerant species, with an average optimal growth temperature of 34°C, followed by *S. paradoxus* and *S. mikatae*, which grow optimally at 33°C and 31°C, respectively; these three species are classified as warm-tolerant. *S. eubayanus, S. kudriavzevii*, and S. arboricola are considered cold-tolerant, with an optimal growth temperature of 26-29°C, showing strong growth limitations at warmer temperatures. *S. uvarum* and *S. jurei* have intermediary thermal tolerance to these two groups, with optimum growth temperatures around 30°C.

**Table 1.** List of *Saccharomyces spp*. yeasts with species names, strain codes, synonymous strain names in public repositories (e.g., the National Collection of Yeast Cultures or Saccharomyces Genome Database), reference genomes used for mapping, and their GenBank or RefSeq assembly ID on NCBI.

| Species name | Strain code | Strain name | Strain for Mapping | Assembly ID |
| --- | --- | --- | --- | --- |
| *S. cerevisiae* | SC1 / SC2 | Y12 / HLJ1 | S288C | GCF_000146045.2 |
| *S. paradoxus* | SP1 / SP2 | YPS138 / IFO1804 | CBS432 | GCF_002079055.1 |
| *S. mikatae* | SM1 / SM2 | NBRC10996 / yHAB336 | IFO1815 | GCF_947241705.1 |
| *S. eubayanus* | SE1 / SE2 | CBS12357 / yHRVM108 | FM1318 | GCF_001298625.1 |
| *S. kudriavzevii* | SK1 / SK2 | NBRC10990 / TJ13M05 | IFO1802 | GCF_947243775.1 |
| *S. uvarum* | SU1 / SU2 | yHAB521 / yHCT77 | CBS7001 | GCA_027557585.1 |
| *S. arboricola* | SA1 / SA2 | TJ14M01 / ZP960 | H-6 | GCA_000292725.1 |
| *S. jurei* | SJ1 / SJ2 | TUM297 / NCYC3947 | UoM1 | GCA_900290405.1<br>(custom gtf) |

All strains used in this study were diploid. Strains were grown from −70°C cryostocks in liquid YPD (1% yeast extract, 2% peptone, 2% glucose) for 24h at 25°C before streaking them on YPD agar (2% agar) for single colonies. Then, a single colony was picked for propagation in 5 mL of liquid YPD for 24h at 25°C.

### Heat wave treatment and serial transfer

We used a temperature of 37°C to simulate the heat waves, which is 3°C higher than the optimal growth temperature of the most thermotolerant species in our experiment (*S. cerevisiae*). All strains were subjected to a repeated heat wave treatment at 37°C to compare their phenotypic and transcriptomic responses. At the same time, all strains were also subjected to a control treatment, where temperature was kept at a constant 25°C (Fig. 1).

**Figure 1.**
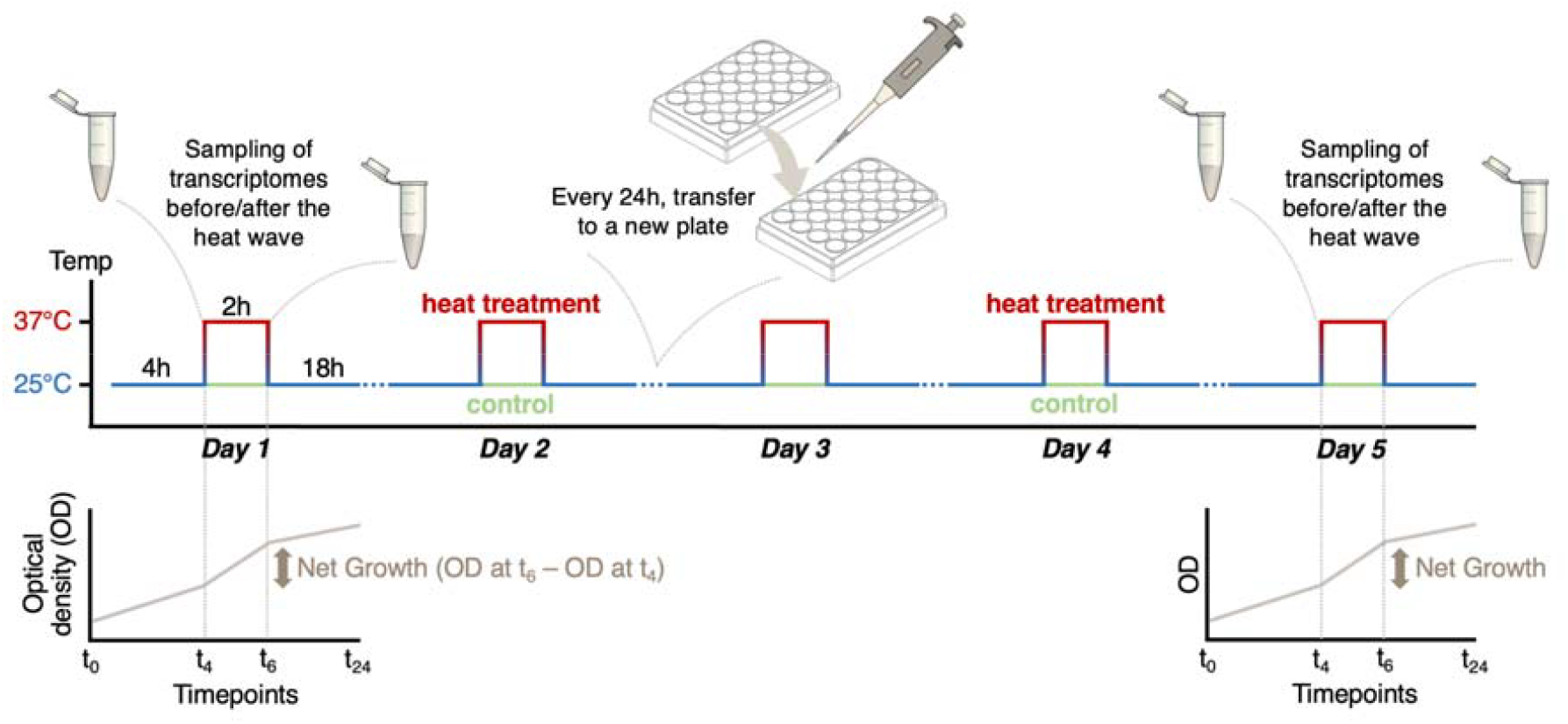
Schematic of the experimental protocol.

Overnight cultures were used to inoculate 24-well flat-bottomed plates, containing 1 mL of YPD per well. We used eight replicated populations per strain, i.e., eight individual wells were inoculated with the same strain and each plate contained populations from two different strains. The two central columns were filled with YPD only (no cells) for spatial separation to avoid cross-contamination. Strains belonging to different species were randomly allocated to plates.

All treated populations were grown for four hours (t_0_ to t_4_) before exposing them to a 2h heat wave at 37°C (t_4_ to t_6_), followed by a recovery period of 18h at 25°C (t_6_ to t_24_). Because populations were in exponential growth phase at t_4_ and the heat wave lasted only two hours (approximately one generation under favorable conditions), changes observed between t_4_ and t_6_ primarily reflect responses to heat stress rather than growth or density-dependent effects. Then, populations were diluted 1:10 using serial transfer into new plates with fresh media. This treatment was repeated for five consecutive days. The number of generations per growth cycle (i.e., per day) was calculated from the OD data using the formula: log2(final_OD_/initial_OD_), for each strain (Supplementary Table S2). The average number of generations over five days per strain was 16.64 ± 0.45 (mean ± sd).

As a control, we used the same serial transfer protocol, but kept populations at 25°C during the heat wave.

### Assessment of heat wave tolerance

Heat wave tolerance was assessed based on the final optical density (OD_600_) of each well, measured with a microplate reader (Sunrise, Tecan) every day at t_0_ (daily transfer into fresh media), t_4_ (*before* the two-hour long heat wave at 37°C, or at 25°C for the control), and t_6_ (*after* the heat wave). On day 1 at t_0_, all strains were normalized to an OD of 0.1.

As a proxy for heat wave tolerance, net growth was calculated during the heat wave period as the difference in OD measured at t_6_ minus the OD measured at t_4_ (ΔOD). Net growth was analysed using linear mixed-effects models with the function lmer from the R package lme4 v1.1-36. Thermo-tolerance group (warm-adapted, thermo-generalist, cold-adapted species; the first term for each effect being the reference level), treatment (control at 25°C or heat wave at 37°C), day (heat wave 1 or 5, i.e., *HW1* or *HW5*), and all two-way and three-way interactions were included as fixed effects. To account for the hierarchical structure of the data, we included random intercepts for species and for strains nested within species, specified respectively as *(1* | *Species)* and *(1* | *Species:Strain)*. This random-effects structure accounts for differences in average growth among species, as well as additional variability among strains belonging to the same species. Variance components were extracted from the fitted model and used to calculate intraclass correlation coefficients, allowing us to quantify the proportion of total variance in net growth attributable to differences among species and among strains within species.

### RNA extraction and sequencing

On day 1 and day 5, a duplicate of each plate was made to collect yeast samples. The transcriptomes were sampled from two replicates of each strain on day 1 and on day 5, *before* and *after* the heat wave (n = 16 strains x 2 days x 2 timepoints x 2 replicates, i.e., 128 samples), by pooling together 500 μL from 4 wells (V = 2 mL per replicate). Samples were then pelleted by centrifugation at 500 x g for 2 min, and the supernatant was carefully removed before immediate flash-freezing in liquid nitrogen and storage at −80 °C.

RNA extraction was performed using YeaStar RNA Kit from Zymo Research. Briefly, 160 μL of YR Lysis buffer and a 5mm stainless steel bead were added to each pellet, and disrupted with QIAGEN TissueLyser II at maximum speed for 1 minute. The following steps were carried out following the manufacturer’s instructions. DNase treatment and RNA Clean-up were then performed using the RNA Clean & Concentrator-5 kit by Zymo Research. Cleaned samples were eluted in 30 μL of DNase/RNase-Free Water. RNA concentration was measured using Qubit with RNA HS Kit (Agilent, Santa Clara, CA, USA), and the quality of seven random samples was assessed using gel electrophoresis. Both reverse-transcription and sequencing were performed by the National Genomics Infrastructure (NGI, Solna, Sweden). Individual libraries were prepared with Illumina TruSeq stranded mRNA (PolyA selection) (Illumina, San Diego, CA, USA), and sequencing was performed on an Illumina Novaseq X 25B flow cell, with 2*150PE. Raw data can be found in the Sequence Read Archive (SRA) database of NCBI under BioProject PRJNA1271760 (reviewer link: https://dataview.ncbi.nlm.nih.gov/object/PRJNA1271760?reviewer=86tdnq8ul63t5mid7v0r7uh8e9).

### Producing transcript annotation file

For all species but *S. jurei*, reference genomes included GTF files with transcript annotations. We therefore produced a custom GTF file for the assembly GCA_900290405.1. Reads from the 16 *S. jurei* samples were first aligned onto the assembly UoM1 FASTA file with HISAT2 v2.2.1 (Zhang *et al*., 2021), and sorted with samtools v1.20 (Li *et al*., 2009). BAM files were then merged together using samtools, before running Braker3 v3.0.8 to annotate the transcripts (Gabriel *et al*., 2024). The GTF file produced was combined with the UoM1 FASTA to extract a FASTA file of the transcripts, using gffread v0.12.7 (Pertea and Pertea, 2020). The completeness of the prediction was assessed by running the transcript FASTA against saccharomycetes_odb10 orthologs using BUSCO v5.4.7 (Manni *et al*., 2021) (2,162 transcripts in total; C:93.9% [S:91.7%, D:2.2%], F:1.5%, M:4.6%, n:2137).

### Transcript quantification and differential expression analysis

Trimming and mapping of the reads were performed following the default parameters from the nf-core/rnaseq pipeline v3.19.0 for each species (Patel *et al*., 2025), using the same assembly for both strains within species (Table 1). For all 128 samples sequenced (16 samples per strain), the proportion of uniquely mapped reads was high (on average 90,79%, minimum 78,56%). The number of transcripts was quantified for over 5000 genes per species, except for *S. arboricola*, totaling around 3,600 genes described.

Read counts by gene were analyzed using the R package DESeq2 v1.46.0 (Love, Huber and Anders, 2014), with *HW1* and *before* as the reference levels, using Salomon’s raw output tables (salmon.merged.gene_counts). A model was built with an interaction between factors Day (*HW1* and *HW5*) and *Heat wave* (*before* and *after*), and *Strain* as an additional effect. Only genes with a minimum of 10 reads in at least two samples per species were kept.

Differential expression was assessed separately for each contrast derived from the DESeq2 model (*before vs after* on HW1, *before vs after* on HW5, HW1 *vs* HW5 *before*, HW1 *vs* HW5 *after*, interaction between heat wave and day effects). For a given contrast, genes were considered as significantly differentially expressed when adjusted p-values were <0.05 and absolute fold change in expression was >2 (over-expression) or <-2 (under-expression), to observe strong, robust, and biologically clear responses. To create gene name correspondence files between *S. cerevisiae* and other *Saccharomyces* species, protein sequences were aligned using BLASTP (Camacho *et al*., 2009), orthologs were identified via OrthoDB (Tegenfeldt *et al*., 2025), and overlapping hits were extracted. Differentially expressed gene names were thus all replaced by *S. cerevisiae* gene names, allowing for comparison between species.

### Transcriptome profiling and functional analysis

Gene counts were normalized and transformed using the variance stabilizing transformation (vst) from DESeq2 before plotting Principal Component Analysis (PCA), using the 500 most variable genes out of the 2,859 common genes. Hierarchical clustering and multidimensional scaling (MDS) were performed to explore global transcriptomic relationships between species and across conditions. Hierarchical clustering was applied to variance-stabilized counts of all expressed genes to assess the phylogenetic structure among the eight *Saccharomyces* species. For MDS, species-averaged expression values were computed for each experimental condition (interaction between day and heat wave conditions). Pairwise Euclidean distances between species were used as input for classical MDS, using the function cmdscale from R/stats v4.4.2 to obtain two-dimensional coordinates.

Gene ontology (GO) terms enrichment analysis was performed on retained differentially expressed genes to identify significantly enriched biological pathways among the gene sets, using the package R/topGO v.2.58.0 (nodeSize 10, algorithm weight01, statistic Fisher). The entire pipeline with the details of the parameters used is available on GitHub (https://github.com/chaberko-lbbe/sacch-rnaseq).

### Conceptual framework for expression dynamics

To interpret the effects of the factors *Day, Heatwave* and their interactions in a biologically meaningful way, we defined a set of predicted responses (Fig. 2), which summarize possible transcriptional dynamics between the first and fifth day of heat waves (HW1 and HW5) and *before/after* each heat wave exposure. Conceptually, our classification of transcriptional responses is adapted from existing frameworks describing gene expression plasticity, acclimation, transcriptional memory, potential transgenerational epigenetic inheritance, and their role in stress tolerance and environmental responses (Chevin, Lande and Mace, 2010; Rivera *et al*., 2021; Li and Zhang, 2026). But because experiments were conducted on clonal populations over a limited number of mitotic generations, all response categories reflect transcriptional changes only, independently from genetic changes.

**Figure 2.**
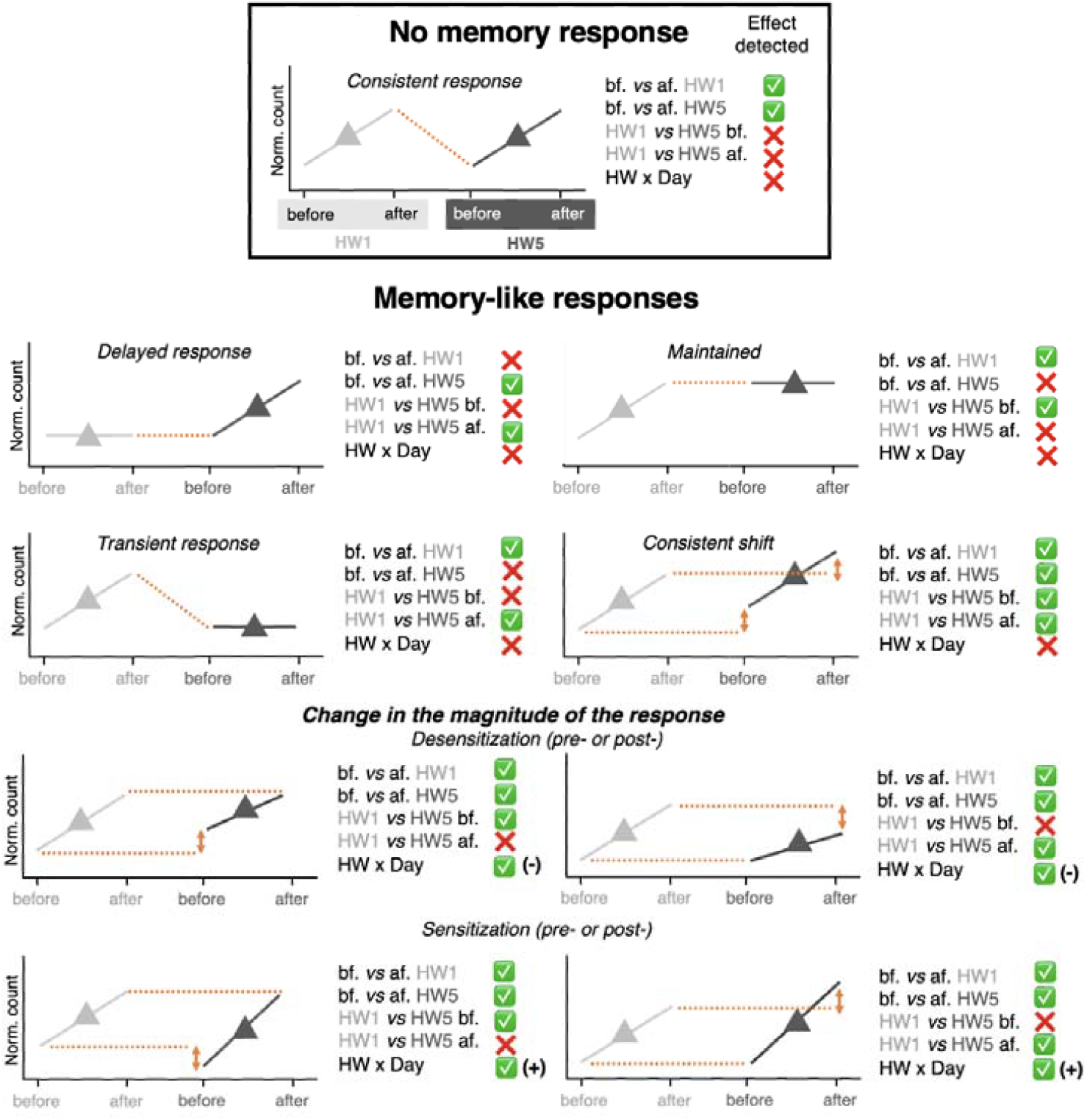
Predictive framework for changes in gene expression when subjected to a heat wave (before and after) across days (HW1 and HW5), including no-memory (consistent) and memory-like responses to heat waves. Triangles indicate mean response per day. Arrows indicate shifts in the response on the fifth day compared to the first, when comparing before (bf.) and/or after (af.) the heat wave across days. Effects predicted to be detected by the model (DeSeq2) are detailed for each case (tick marks and crosses). Only positive changes are shown here (i.e., over-expression), but negative changes (i.e., under-expression) follow the same patterns.

Predicted changes in gene expression showing *no-memory responses* are described as *consistent* changes across events (i.e., differentially expressed genes upon heat wave, shared in both HW1 and HW5). Conversely, we interpreted genes showing differences in their response during the first and fifth heat wave as *memory-like* responses. These are operational expression categories, not direct evidence of epigenetic mechanisms.

*Memory-like* responses encompassed *transient* expression (i.e., differentially expressed genes unique to HW1), *maintained* expression (i.e. genes consistently expressed in HW5 to the level reached after HW1), *delayed* expression (i.e., differentially expressed genes unique to HW5), or genes with a *shift* in expression *before* and/or *after* the fifth heat wave compared to the first heat wave. We further subdivided shifts into *consistent shifts* (i.e., shift in the same direction *before* and *after* the heat wave), or *change in the magnitude of the response* (i.e., shift either only pre- or only post-the heat wave). When showing a change of magnitude, the intensity of the response (slope) can be either reduced (*desensitization*) or increased (*sensitization*). This conceptual framework guided our biological interpretations of DESeq2 outputs and the classification of expression profiles across conditions. For each prediction of changes in expression, we defined a set of conditions where the effect was expected to be present (green tick marks in Fig 2) and absent (red crosses in Fig 2). GO terms identified as significantly enriched for each effect detected were then classified into response categories based on their presence or absence across these predicted effects.

## Results

### Species natural thermotolerance impacts growth response to heat waves

Average net growth of populations was measured during the two-hour heat wave at 37°C, as well as for the control staying at 25°C during the same period, for both day 1 and 5 (Fig. 3). We categorized species into three thermo-tolerance groups following previous work: we considered populations of *S. cerevisiae, S. paradoxus* and *S. mikatae as warm-adapted, S. jurei* and *S. uvarum* as thermo-generalist, and *S. arboricola, S. kudriavzevii* and *S. eubayanus* as cold-adapted (Molinet and Stelkens, 2025).

**Figure 3.**
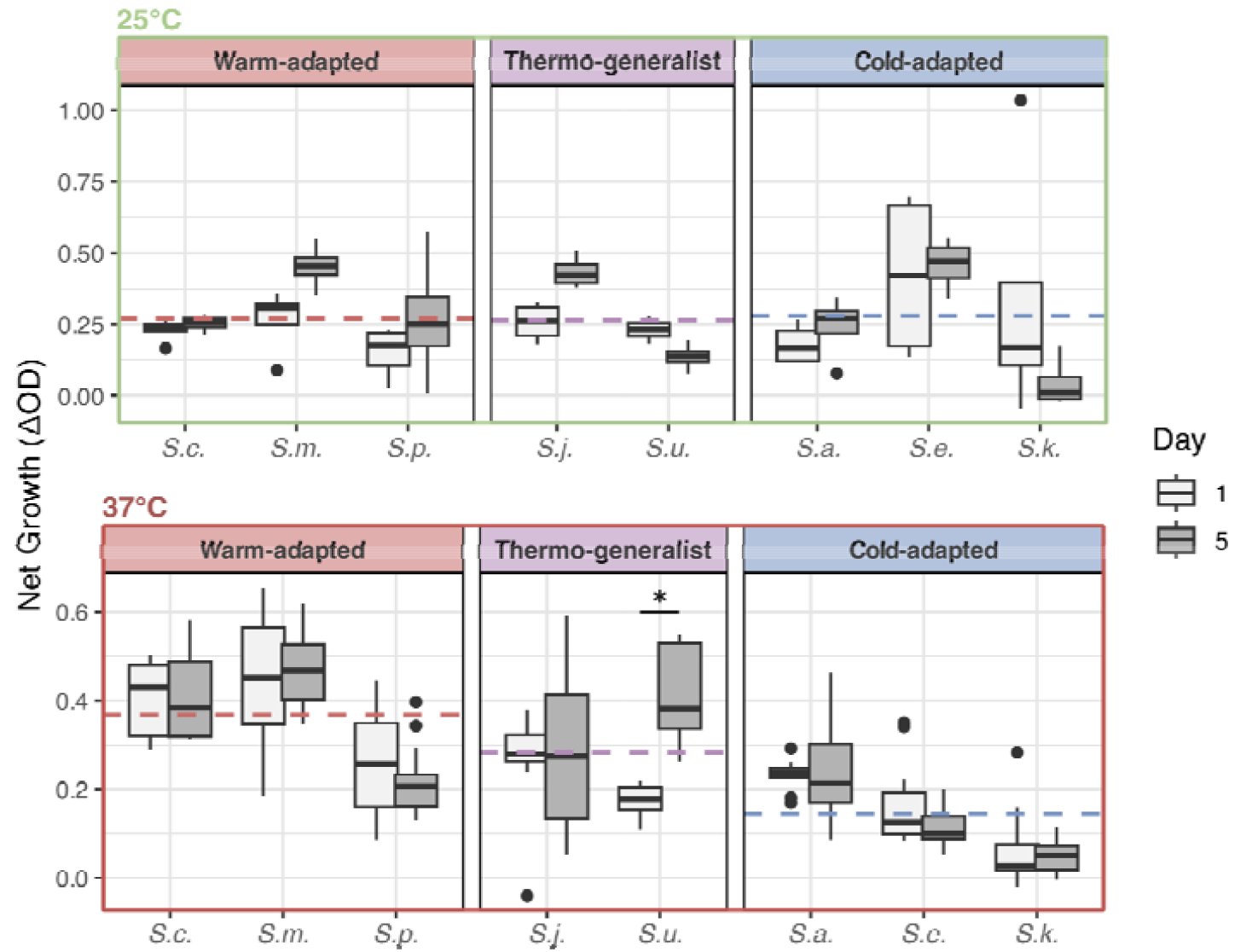
Net growth during consecutive heat waves on day 1 and day 5. Differences in optical density (*after* minus *before* the heat wave, ΔOD) averaged across sixteen replicates per *Saccharomyces* species (two strains per species, eight replicates per strain). Asterisks indicate significant differences between days (Wilcoxon test with Bonferroni correction for multiple testing (*, p < 0.05). Dashed lines represent mean values for species groups considered warm-adapted, thermo-generalist, and cold-adapted.

We analysed net growth using linear mixed-effects models accounting for both experimental factors and hierarchical biological structure. *Species* accounted for 16.7% of the total variance in net growth, while *Strain* within species explained an additional 10.1%, suggesting that differences in growth responses are more pronounced among species than among strains belonging to the same species. For the main effects, net growth was significantly higher in the heat wave treatment than in the control treatment (estimate = 0.16 ± 0.06 SE, p = 0.007). We also found a temporal positive effect, with net growth increasing significantly between day 1 and day 5 (estimate = 0.03 ± 0.01 SE, p = 0.045). No significant main effect was found between thermo-tolerance groups, but significant interaction effects indicate that responses to heat waves differed among them. Indeed, compared to warm-adapted species, thermo-generalist and cold-adapted species showed a significantly smaller increase in net growth under repeated heat waves relative to the control condition (thermo-generalist: estimate = −0.195 ± 0.090 SE, p = 0.031; cold-adapted: estimate = −0.324 ± 0.081 SE, p = 7.68e-05). This pattern is reflected in the group means: under the heat-wave treatment, warm-adapted species showed the highest net growth (mean ± sd: 0.36 ± 0.11), followed by thermo-generalist species (0.28 ± 0.12), whereas cold-adapted species showing the lowest growth (0.14 ± 0.09; Fig. 3). In contrast, under the control treatment at 25°C, net growth was similar among all species groups (warm-adapted: 0.27 ± 0.14; thermo-generalist: 0.27 ± 0.12; cold-adapted: 0.28 ± 0.26). Overall, the heat wave treatment resulted in a +0.09 increase in mean net growth in warm-adapted species, little change in thermo-generalist species (+0.01), and a decrease of −0.14 in cold-adapted species relative to the control treatment, consistent with the significant *treatment* × *thermo-tolerance group* interaction detected by the mixed-effects model.

A significant interaction between cold-adapted species and *Day* further suggests that net growth decreased more strongly from day 1 to day 5 in this group than in warm-adapted species (estimate = −0.043 ± 0.019 SE, p = 0.027). This pattern is consistent with a relative decline in growth performance of cold-adapted species across successive heat waves. No significant interaction between *Treatment* and *Day* or any three-way interactions were detected.

Within species, net growth during heat waves was similar between the first and last heat wave for all species except for the thermo-generalist *S. uvarum*, which showed significantly higher growth on day 5 than on day 1 (Wilcoxon test with Bonferroni correction, p = 5.92e-06). Under control conditions, no species showed a significant difference in net growth between day 1 and day 5.

### Differences in transcriptomic response to heat waves between species

RNA-sequencing was performed to compare the transcriptomes of the eight species with two strains each *before* and *after* the heat wave, on day 1 (*HW1*) and day 5 (*HW5*). The transcriptomes of each species were first compared by principal component analysis (PCA; Fig. 4). Transcriptomes were separated into four clusters along the two first component axes: ‘*before*-day 1’ (light grey triangles), ‘*before*-day 5’ (dark grey triangles), ‘*after*-day 1’ (light grey circle) and ‘*after*-day 5’ (dark grey circle), except for *S. jurei* and *S. arboricola*. For the last two species, the primary source of variation (PC1, explaining 60% and 92% of the variance, respectively) clearly corresponded to the heat wave effect (*before* vs *after*), while PC2 did not clearly separate day 1 from day 5, and only explained a small portion of variance (18% for *S. jurei*, and only 3% for *S. arboricola*).

**Figure 4.**
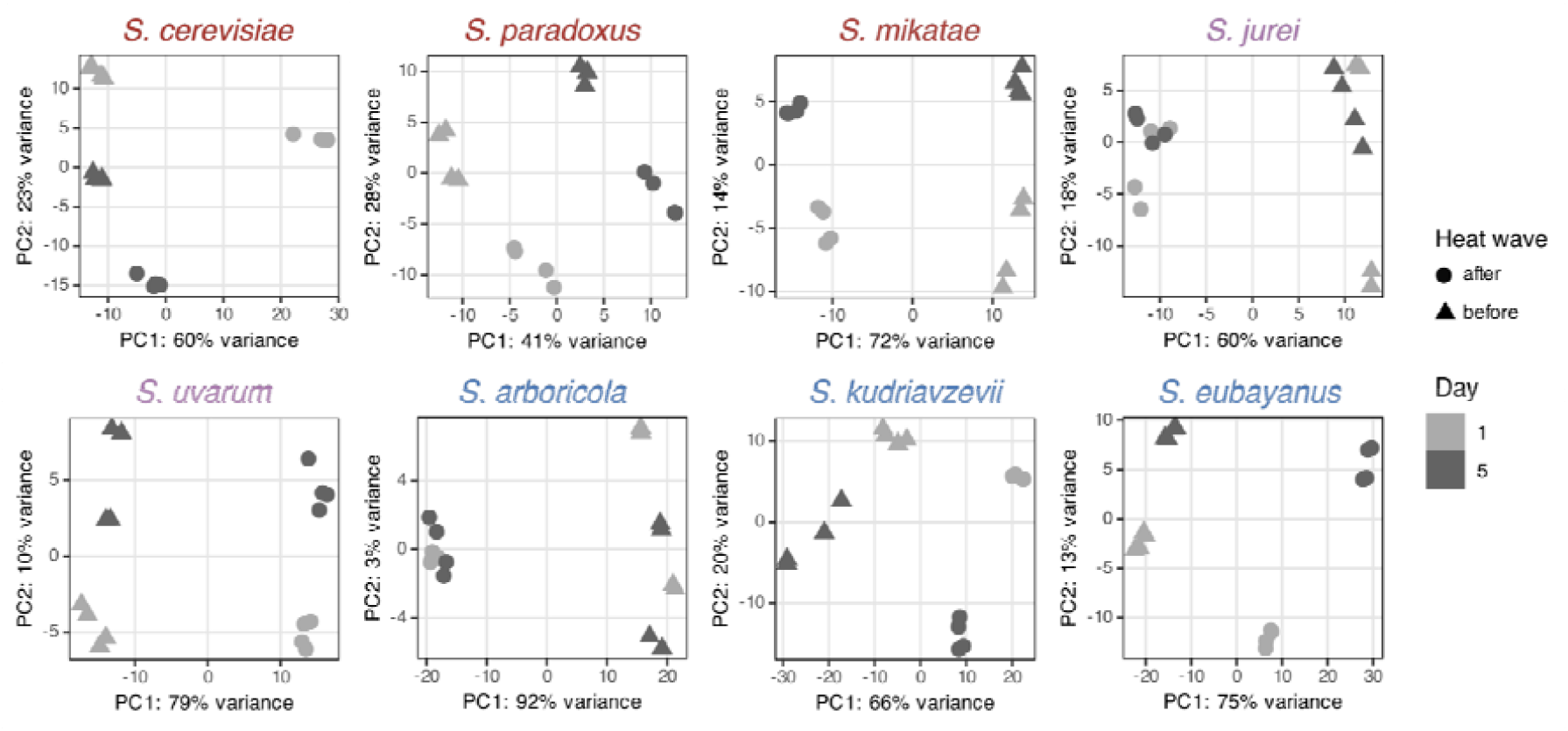
Projection of all *Saccharomyces* transcriptomes on the first two principal components using PCA. Data points represent the top 500 most variable genes of a sample (default parameter for plotPCA function, out of 2,859 common genes) and are shaded by day 1 or 5. The shape of the points shows *before* or *after* the heat wave. Every point therefore represents a replicate within a strain for a combination of conditions (n = 16 per species, 8 per strains, 4 per condition, 2 replicates). Read counts for genes were obtained from DESeq2 and processed with *vst*, which computes a variance stabilizing transformation.

Overall, the heat wave effect explained the largest amount of variance in the transcriptomes of all species, followed by the effect of the day sampled, which had variable impact depending on the species. Strains were closely clustered within species depending on experimental conditions (Supplementary Fig. S1).

Hierarchical clustering of species on their global expression profiles revealed a clear phylogenetic structure (Fig. 5A). Warm-adapted *S. cerevisiae, S. paradoxus* and *S. mikatae* clustered with the thermo-generalist *S. jurei*, while the cold-adapted species *S. kudriavzevii, S. arboricola* and *S. eubayanus* were grouped with the thermo-generalist *S. uvarum*. To examine whether these relationships were consistent across sampling time-points, we performed multidimensional scaling (MDS) using species-averaged expression distances, separating the four time points: ‘*before*-day 1’, ‘*before*-day 5’, ‘*after*-day 1’ and ‘*after*-day 5’ (Fig. 5B).

**Figure 5.**
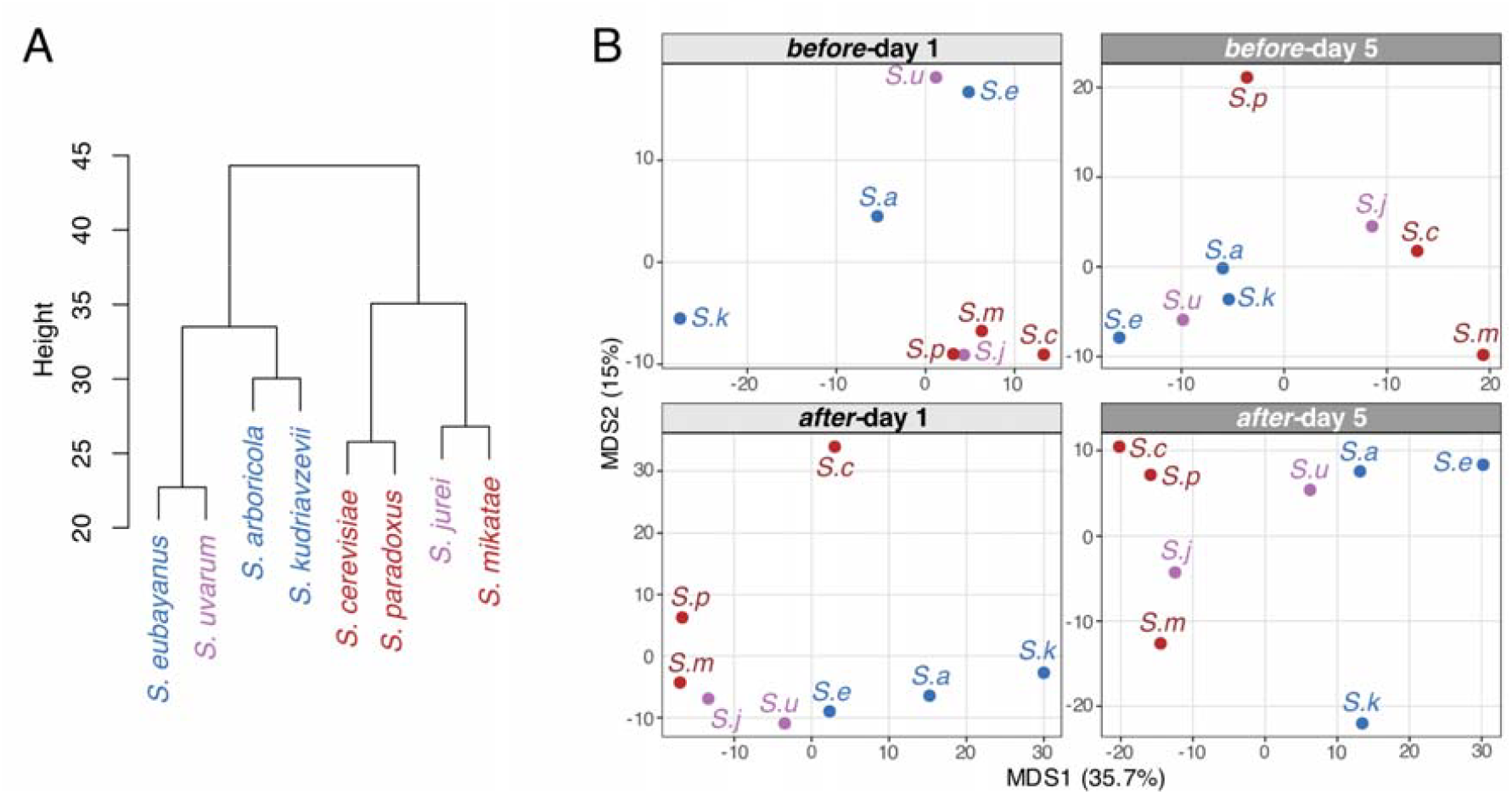
Hierarchical clustering and multidimensional scaling (MDS) of *Saccharomyces* species based on expression profiles. **A)** Dendrogram of species relationships derived from hierarchical clustering of Euclidean distances between species-averaged expression profiles across conditions. **B)** MDS plots of species similarity under four sampling time-points: *before-*day 1, *before-*day 5, *after*-day 1 and *after*-day 5. Species are colored by thermotolerance, following Molinet and Stelkens (2025).

The first two MDS axes explained 35.7% and 15.0% of the variance, respectively, capturing just over half of the overall distance structure. Before heat wave exposure, the three warm-adapted species and *S. jurei* clustered closely at the ‘*before*-day 1’ time point, whereas the three cold-adapted species and *S. uvarum* clustered together at the ‘*before*-day 5’ time point,

After heat wave exposure, the clustering patterns shifted. At the ‘*after*-day 1’ time point, *S. cerevisiae* was separated from the other species along MDS2, while MDS1 broadly distinguished warm-adapted, thermo-generalist, and cold-adapted species. By ‘after-day 5’, *S. cerevisiae* grouped with the other warm-adapted species on MDS1, consistent with the hierarchical clustering (Fig. 5A).

Overall, both hierarchical clustering and MDS consistently revealed two major groups that broadly corresponded to species with higher and lower heat-wave tolerance: (i) *S. cerevisiae, S. paradoxus, S. mikatae and S. jurei*, and (ii) *S. eubayanus, S. uvarum, S. arboricola* and *S. kudriavzevii*. The two thermo-generalist species (*S. uvarum* and *S. jurei*) clustered with opposite groups indicating that thermal niche breadth alone does not lead to transcriptional similarity. These patterns suggest that species with similar thermal adaptation also tend to exhibit more similar global transcriptional responses to repeated heat wave exposure.

To further investigate the molecular basis of the observed transcriptomic changes, we performed differential gene expression analysis. Building on the PCA results, we focused on identifying genes whose expression was significantly affected by the heat wave treatment (*before* and *after*), sampling day (day 1 and day 5), and their interaction, while accounting for strain-specific variation. We identified between 0 and 1061 significantly differentially over- or under-expressed genes, depending on the species and conditions (Supplementary Table S3). For each set of differentially expressed genes, enriched Gene Ontology terms were identified within the Biological Process categories (Supplementary Table S4). Only one GO term was detected as enriched for genes differentially expressed for the condition Strain (siderophore transport, under-expressed in *S. kudriavzevii)*. This condition was therefore not further analysed.

### Transcriptomic response to the first heat wave

The first heat wave can be considered a single heat shock. To analyse how species responded, *before* and *after* samples from *day 1* were compared (**HW1** effect). Upon HW1, over 200 genes were detected as over-expressed for each species (786 for *S. cerevisiae*, 239 for *S. paradoxus*, 338 for *S. mikatae*, 342 for *S. jurei*, 575 for *S. uvarum*, 482 for *S. arboricola*, 615 for *S. kudriavzevii* and 658 for *S. eubayanus*), and over 40 genes as under-expressed (919 for *S. cerevisiae*, 41 for *S. paradoxus*, 257 for *S. mikatae*, 181 for *S. jurei*, 364 for *S. uvarum*, 366 for *S. arboricola*, 584 for *S. kudriavzevii* and 421 for *S. eubayanus*) (Supplementary Table S3). Multiple heat shock proteins (labelled on Fig. 6) were detected as over-expressed in almost all species, such as *HSP82*, a negative regulator of the Hsf1p-dependent heat shock response, or *HSP26*, which has a chaperone activity. Other kinds of chaperones were found to be under-expressed (e.g., in *S. cerevisiae, S. eubayanus, S. kudriavzevii*), for instance cytoplasmic ATPases that are ribosome-associated molecular chaperones, such as *SBB1* or *SSB2* (labelled on Fig. 6).

**Figure 6.**
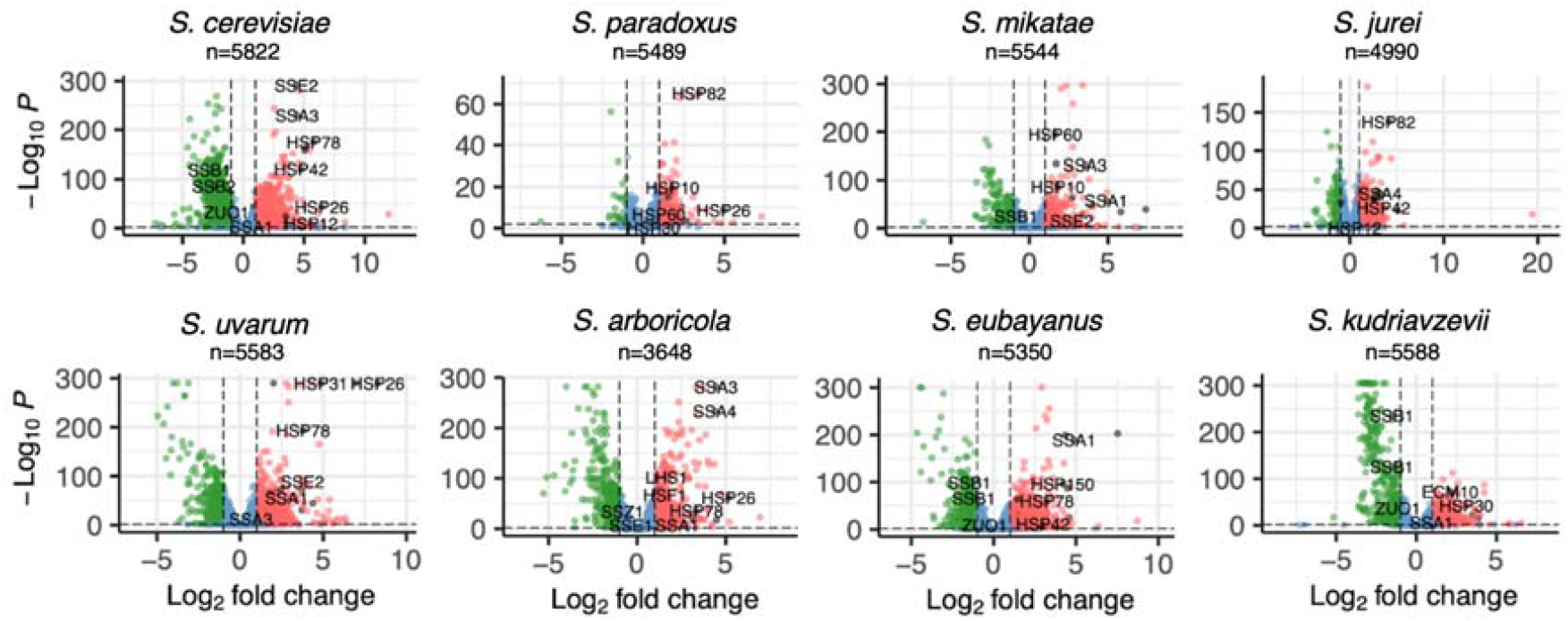
Volcano plot on heat wave effect (*before* vs *after*) during *day 1* (⍰⍰p_adj_ < 0.05, LFC >⍰1 or <-1, represented by dashed lines). Heat shock proteins are labelled (manual database built with Uniprot).

### Changes in transcriptomic response across daily heat waves

To interpret changes in transcriptomic responses across repeated heat waves, we applied the conceptual framework we described above (Fig. 2). Out of 429 functional enrichment patterns, 67 were assigned to either no-memory or memory-like transcriptional responses categories (Supplementary Table S5), with 49 no-memory responses and 18 memory-like responses (Fig. 7).

**Figure 7.**
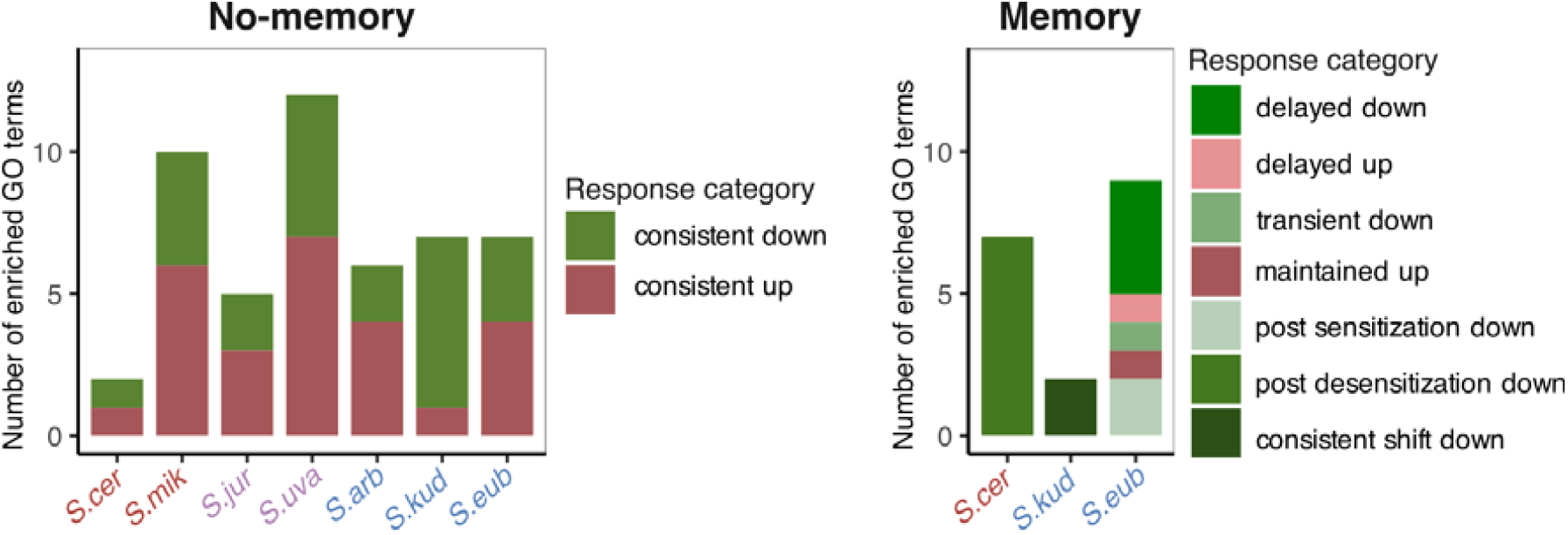
Number of enriched gene ontology (GO) terms associated with no-memory (left) and memory-like (right) transcriptional response categories across yeast species exposed to repeated heat waves.

### Consistent no-memory responses across heat waves

We first focused on GO terms associated with no-memory responses (Fig. 2), i.e., consistent responses across heat waves. Both up-regulated genes (11 GO terms enhanced) and down-regulated genes (14 GO terms repressed) were detected as triggered similarly by repeated exposure across all species (Fig. 8).

**Figure 8.**
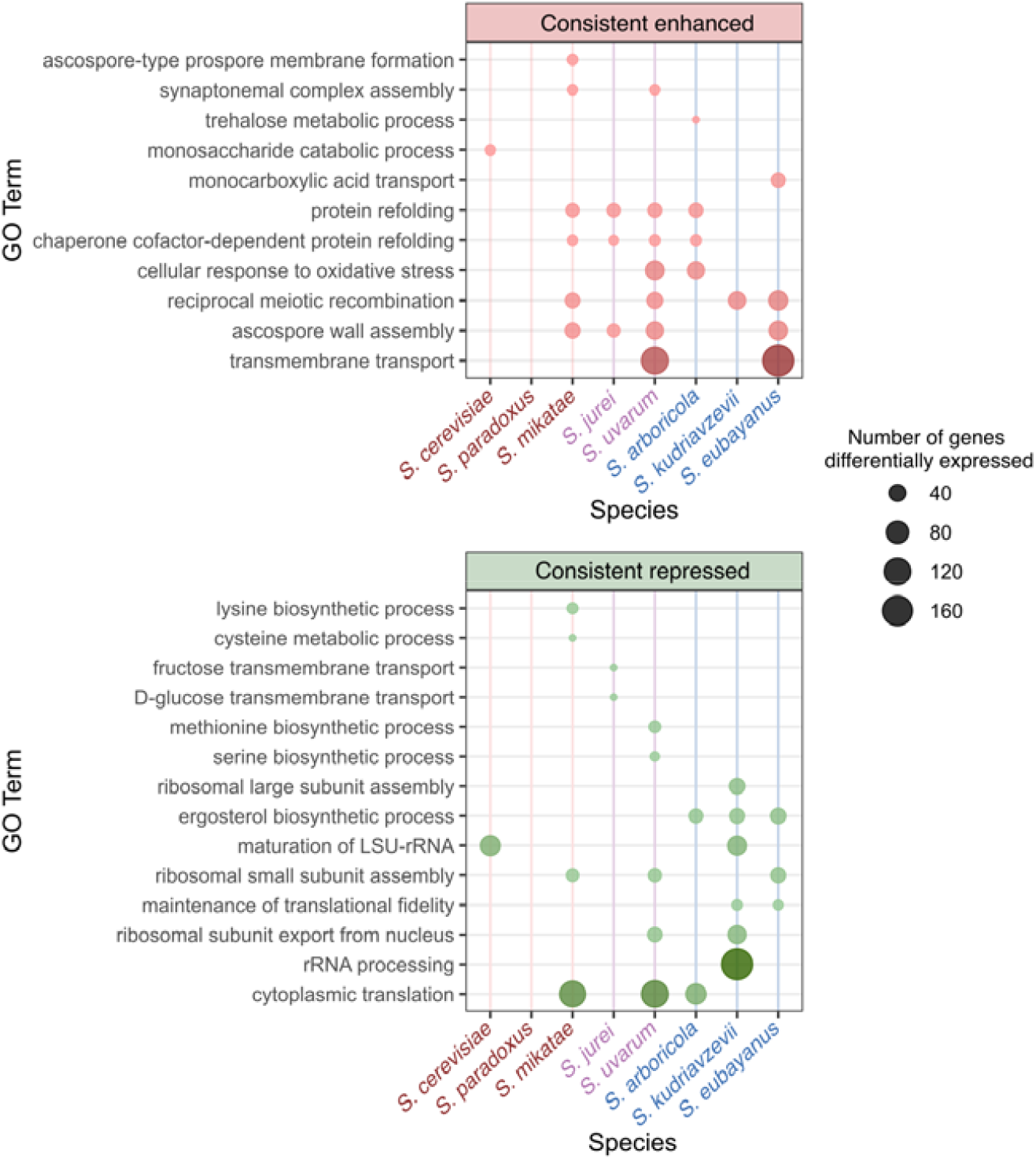
Consistently enriched GO terms grouped according to the conceptual framework of potential heat wave responses (Fig. 2). GO terms significantly enriched among up-regulated (enhanced) or down-regulated (repressed) genes upon both the first and fifth heat wave (HW1 and HW5) were identified for each *Saccharomyces* species, and subsequently grouped into broader functional categories derived from the conceptual framework. Species names are colored according to their thermotolerance (warm, generalist, or cold). Dot size indicates the total number of differentially expressed genes annotated to each GO term (sum across HW1 and HW5).

Among the consistently enhanced GO terms, several meiosis-related processes (*ascospore-type prospore membrane formation, ascospore wall assembly, synaptonemal complex assembly* and *reciprocal meiotic recombinations*) were induced, in warm-adapted (*S. mikatae, S. paradoxus*), thermo-generalist (*S. jurei, S. uvarum*) and cold-adapted species (*S. kudriavzevii, S. eubayanus*). This suggests a broadly shared activation of sporulation-associated transcriptional programs in response to both heat waves across all species. Similarly, enrichment for protein folding and refolding terms was seen in species from diverse thermal tolerance profile (*S. mikatae, S. jurei, S. uvarum*, and *S. arboricola*), highlighting a broad requirement for chaperone-mediated proteostasis. In contrast, *cellular response to oxidative stress* and *transmembrane transport*, including proteins from the Hsp70 family, were found to be shared only by cold-adapted and one generalist species (*S. uvarum* and *S. arboricola* or *S. eubayanus*). Finally, expression processes associated with different sugars were species-specific, such as trehalose for *S. arboricola*, a sugar known to be stress protectant (Zhang, Zhang and Li, 2020).

For down-regulated genes, we found many genes associated with ribosome biogenesis and translation pathways, translation being the ribosome-mediated process by which mRNA is translated into proteins in the cytoplasm. The repression of translation via *cytoplasmic translation or maintenance of translational fidelity* was detected in species from diverse thermal tolerance profile (*S. mikatae, S*.*uvarum, S. arboricola, S. kudriavzevii* and *S. eubayanus*). Ribosome-related pathways followed the same trend, underscoring a conserved repression of protein synthesis upon repeated heat waves. We found five different pathways related to ribosome biogenesis in almost all species: maturation of *LSU-rRNA, ribosomal small* and *large subunit assembly, ribosomal subunit export from nucleus*, and *rRNA processing* (*S. cerevisiae, S. mikatae, S*.*uvarum, S. arboricola, S. kudriavzevii, S. eubayanus*). Beyond ribosomal repression, several transcriptome cluster-specific metabolic processes were also repressed. For instance, the cold-adapted *S. kudriavzevii, S. arboricola* and *S. eubayanus* species showed enrichment for *ergosterol biosynthetic process*, while pathways related to sugar transport such as *D-glucose, fructose*, and *mannose transmembrane transport* were only detected in the thermogeneralist *S. jurei*. Species-specific enrichment in genes encoding amino acids biosynthesis processes were also observed - lysine and cysteine for *S. mikatae*, methionine and serine for *S. uvarum*.

In summary, repeated heat wave exposure revealed a broadly shared set of consistent transcriptomic responses across species, including the activation of meiosis-related processes and facilitating protein folding, and the repression protein translation overall.

### Most memory responses involved changes in response magnitude

Next, we explored GO terms predicted to show memory-like responses (Fig. 2), encompassing genes with differences in the expression levels observed during the first and fifth heat wave (*maintained, transient, delayed*), or showing a *shift* in response.

Only one GO term, related to glycolytic processes, showed a transient repression in the cold-adapted *S. eubayanus*, suggesting that most functional pathways were not rapidly reversed after the first heat exposure (Fig. 7). *S. eubayanus* also showed some delayed enhanced responses, where genes involved in glucan catabolic processes became over-expressed only after the final heat wave: three GO terms involved in ribosome biogenesis and one in RNA methylation showed delayed repression in *S. eubayanus*. We found only one maintained response, for the GO term protein refolding in *S. eubayanus*, which was detected as over-expressed upon the first heat wave and had a significantly higher expression *before* the last heat wave when compared to *before* the first. Similarly, we observed the genes of only two GO terms as showing a consistent shift pattern, with an under-expression upon both heat waves that became more intense during the last heat wave. These two GO terms, related to translation, were detected in *S. kudriavzevii*.

Multiple memory-like responses were consistent with our prediction for (de)sensitization, with seven GO terms associated with under-expressed genes showing post-desensitization in *S. cerevisiae*, and two GO terms associated with under-expressed genes showing post-sensitization in *S. eubayanus* (i.e., 9 out of 18). Of these GO terms, we found three were involved in ribosome biogenesis and four in translation for S. cerevisiae, and one each in ribosome biogenesis and translation for *S. eubayanus*. Interestingly, two of the GO terms were found to be associated with memory-like responses in both the cold-adapted *S. eubayanus* and the warm-adapted *S. cerevisiae* (‘maturation of SSU-rRNA from tricistronic rRNA transcript’, GO:0000462, and cytoplasmic translation, GO:0002181). These two species represent thermal tolerances at the opposite extremes in the *Saccharomyces* genus. Although both species expressed ribosomal genes (in GO:0000462) at similar levels before the first and fifth heat wave, and under-expressed them after the heatwave on both days, the magnitude and direction of these species’ responses differed: in *S. cerevisiae*, these ribosomal genes were more strongly under-expressed after the first heat wave than after the fifth heat wave, whereas in *S. eubayanus*, we found the opposite pattern (Fig. 9). Thus, memory-like modulation was concentrated in ribosome biogenesis and translation, suggesting that repeated heat waves altered the magnitude of a shared core response rather than broadly rewiring the transcriptome.

**Figure 9.**
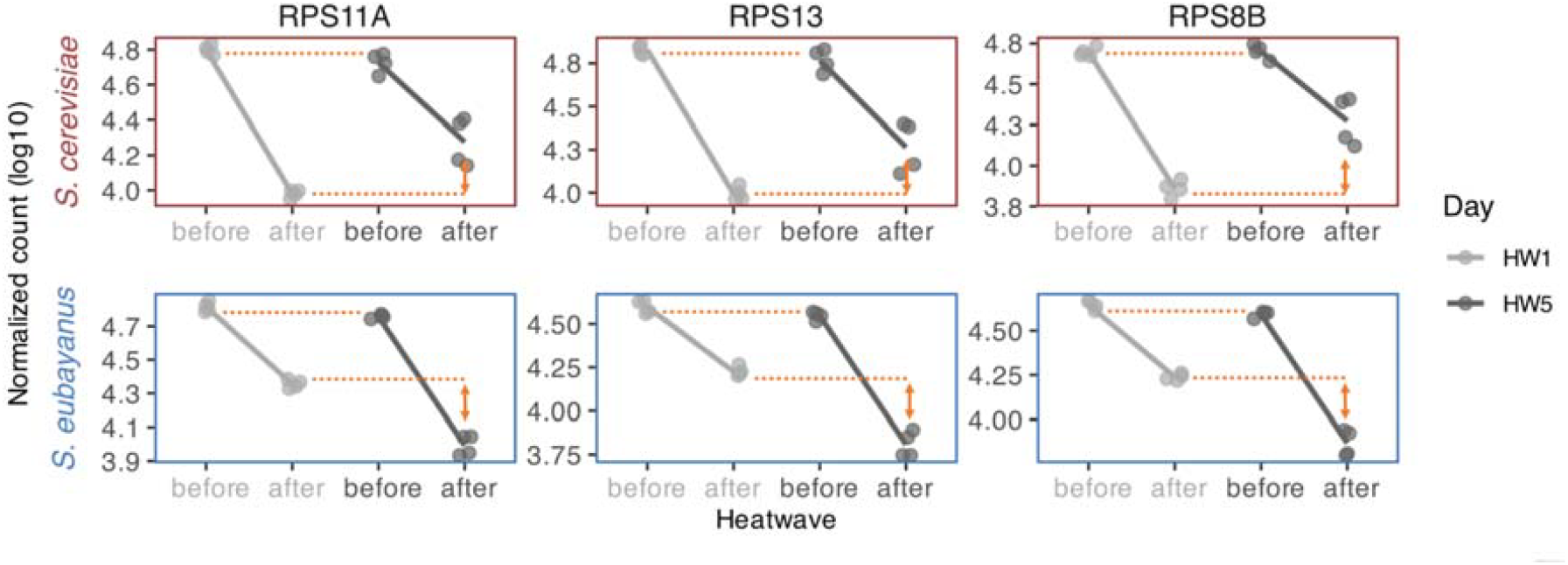
Expression of genes at different sampling time points *before* and *after* exposure to repeated heatwaves. Normalized transcript counts (with log10-transformation of the y-axis) are shown for individual samples (4 replicates per species), *before* and *after* the heat wave on day 1 (HW1) and day 5 (HW5). Lines indicate mean expression. Panels are labeled by the top 3 genes (in read counts) detected in both species from the ribosomal GO term GO:0000462, which follow the prediction of “post-desensitization” for *S. cerevisiae*, and “post-sensitization” for *S*.*eubayanus)*.

## Discussion

Understanding how organisms cope with heat waves is essential for predicting organismal responses to climate change, especially given the fact that heat waves are likely becoming more frequent and intense in the future (Domeisen *et al*., 2023). Here, we show that thermal tolerance fundamentally shapes physiological and transcriptomic responses to repeated heat waves across the *Saccharomyces* yeast genus. We uncovered patterns associated with species-level thermal niches (represented here by two genetically different strains belonging to eight species in total), suggesting that thermal tolerance is a key predictor of species-level responses to recurring heat stress. Warm-adapted species maintained the highest growth rates across heat exposures, whereas cold-adapted species grew less well during the heat wave and showed a relative decline in growth across successive exposures. Thermo-generalists displayed intermediate patterns, with one species improving growth after repeated heat waves, suggesting potential for partial acclimation. Under control conditions at 25°C, all groups maintained similar growth, highlighting that these patterns emerge specifically under heat stress. Convergence in a core heat-stress program that is conserved across species included the induction of proteostasis and repression of translation and ribosome biogenesis. Memory-like transcriptional adjustments, i.e., modulations of gene expression by prior exposure, were species-specific. Although the two strains sampled per species showed similar responses under our experimental conditions (Supplementary Fig. S1), they cannot capture the full extent of intraspecific diversity. Future studies including a broader sampling of natural isolates will therefore be needed to determine how broadly these species-level patterns generalize. Together, these findings suggest distinct evolutionary strategies for coping with the thermal stress exerted by heat waves, and provide a comparative perspective on species resilience under ongoing climate warming.

### Thermal niche predicts phenotypic performance under heat waves

Our growth and transcriptome dynamics suggest three complementary strategies for coping with heat waves. First, warm-adapted species showed resistance, characterized by high performance under heat and the capacity to limit growth loss across repeated events (Fig. 3). Second, cold-adapted species showed relative resistance, i.e., they suffered an immediate performance cost during heat exposure but maintained stable, reduced growth across repeated heat waves. Third, one of the thermo-generalists showed phenotypic acclimation: *S. uvarum* grew better after successive heat waves. However, we only detected enriched GO terms that were consistently enhanced or repressed for this species (Fig. 7, Fig. 8), and no memory-like responses. This suggests that *S. uvarum*’s acclimation capacity may rely on regulatory mechanisms beyond changes in gene expression, such as post-transcriptional or metabolic adjustments. Evidence from other systems shows that heat acclimation can involve persistent changes in the metabolome even after transcriptomic responses return toward baseline (Garcia-Molina *et al*., 2020) and that heat stress can induce extensive alternative splicing, underscoring the role of post-transcriptional regulation (Chen and Li, 2017).

A key question raised by these patterns is whether they reflect intrinsic regulatory strategies, differences in thermal mismatch, or both. Molinet & Stelkens (2025) showed that adaptive shifts in thermal performance curves across *Saccharomyces* species follow distinct evolutionary trajectories and involve trade-offs between upper thermal limits and maximal growth. Thus, the impact of a given temperature depends on where it falls relative to a species’ thermal optimum. This distinction is directly relevant to our experimental design, using a standardized heat wave regime (37 °C). For warm-adapted species, 37 °C represents a moderate deviation from their optimal temperature (+3-4°C), whereas for cold-adapted species it constitutes a much stronger thermal challenge, exceeding their optima by more than 10 °C (Supplementary Table S1). This variation in the relative severity of heat stress likely contributes to differences in growth performance, which in turn may influence gene expression, given the close association between growth and transcriptomic state in yeast (Pelechano and Pérez-Ortín, 2010). Therefore, we cannot exclude that part of the observed variation reflects indirect effects of species-specific growth and physiological states under heat stress. However, our analysis does not aim to compare the overall intensity of heat wave responses among species. Instead, we focus on within-species transcriptional changes occurring *before* and *after* heat wave events across days, rather than comparing expression states between control and heat wave-temperatures. This approach allowed us to capture the dynamics and magnitude of transcriptional responses to heat-wave events, while reducing the confounding effects of interspecific physiological differences.

Quantifying the ecological impact of a heat wave is inherently complex, as it integrates multiple interacting dimensions, including temperature intensity, duration, frequency, and the microclimatic conditions experienced by organisms. In natural environments, thermal exposure can deviate substantially from ambient air temperatures. For instance, substrates such as sun-exposed tree bark, where *Saccharomyces* yeasts are often found (Mozzachiodi *et al*., 2022; Pinto *et al*., 2025), can reach surface temperatures several degrees above surrounding air, generating short but intense thermal extremes that are rarely captured by standard laboratory conditions (Sheppard, Morhart and Spiecker, 2016). Future experiments could incorporate microclimate data and ecological niche modelling to better approximate the natural variability in both the magnitude and duration of heat waves.

### Convergence in core responses to heat wave stress

Across species with different thermal tolerance, we observed convergence of increased expression of proteostasis pathways and repression of translation and ribosome biogenesis after the first and upon the last heat wave (Fig. 8). These patterns mirror the well-described yeast heat-shock response, driven largely by Hsf1, with strong up-regulation of Hsp70/Hsp90 chaperones and global translational down-regulation through TOR inhibition and ribosome biogenesis shutdown (Jacob *et al*., 2012; Shalgi *et al*., 2013; Mühlhofer *et al*., 2019). Ribosome repression is a typical response to heat and proteotoxic stress across fungi and other eukaryotes, such as plants (Merret *et al*., 2017) or mammals (Shalgi *et al*., 2013), acting to reduce protein load and protect proteostasis.

Meiosis- and sporulation-associated gene sets were also consistently up-regulated across heat waves in most species. The induction of sporulation-associated genes likely reflects activation of early, stress-responsive components of the meiotic program rather than full meiotic entry. Yeast can temporarily upregulate early sporulation transcription modules in the event of nutritional limitation and other stresses without completing meiosis, which enables cells to survive longer or prepare for quiescence (Neiman, 2011).

Fay *et al*. (2023) demonstrated that species of *Saccharomyces* display divergent heat-shock transcriptional responses that correlate with their thermal tolerance, with stronger induction of protein-folding pathways and greater repression of translation-related genes in cold-tolerant species (*S. kudriavzevii* and *S. uvarum*) relative to warm-tolerant species (*S. cerevisiae* and *S. paradoxus*). In line with this, we also found these pathways to be consistently enhanced and repressed in *S. uvarum*, but not in *S. cerevisiae* and *S. paradoxus*. However, because our design included several heat waves, this allows us to distinguish between responses that remained stable across exposures from those that changed. Here, we show for the first time that pathways repressed only in cold-adapted species, such as translation, were also repressed in warm-adapted species, but with changes in magnitude depending on the day (first heat wave *vs* last). This indicates that besides sharing a common repertoire of consistent responses comparable to a heat shock, different species of *Saccharomyces* rely on divergent strategies in modulating their responses to repeated heat waves.

### Divergent memory-like transcriptional responses to heat waves

In contrast to the broadly conserved, immediate heat wave response, memory-like transcriptional response was limited (Fig. 7) and predominantly observed in ribosome biogenesis and translation pathways. Notably, these changes manifested in opposite directions in the most warm-adapted species (*S. cerevisiae*) and the most cold-adapted species (*S. eubayanus*), suggesting that memory-like responses following repeated heat wave exposure are highly species-specific. Consistent with previous studies, transcriptional memory generally represented a more subtle modulation of the initial stress response than a complete reprogramming of gene expression, often resulting in a “dampened” response upon repeated heat exposure (Serrano-Quílez *et al*., 2025). These divergent strategies may reflect distinct energy-allocation constraints. In *S. cerevisiae*, desensitization reduces the cost of sustaining stress defenses once initial protection is established. Sensitization on the other hand, as observed in *S. eubayanus*, may indicate either cumulative damage on ribosome assembly, due to a higher stress load, and requiring increasingly strong repression as a reaction; or a compensatory mechanism for maintaining growth (see Figure 3, no significant difference in the net growth of *S. eubayanus* under heat wave treatment on day 1 and 5).

An important observation from our analysis is that memory-like behavior was restricted to a limited set of pathways, rather than being broadly distributed across the transcriptome. This suggests that mechanisms other than the simple re-activation of transcription factors are involved. Ribosome biogenesis and translation are known to be rapidly remodelled during environmental stress as part of the well-described Environmental Stress Response, where these energetically expensive processes redirect cellular resources away from growth and towards stress protection (Gasch, 2026).

How repeated heat-wave exposure generates these persistent, pathway-specific transcriptional responses remains an open question. In multicellular eukaryotes, repeated environmental stress can induce persistent regulatory changes that modulate subsequent transcriptional responses through diverse epigenetic mechanisms (Vihervaara *et al*., 2021; Tehrani *et al*., 2025). Such transcriptional memory has been associated with incomplete resetting of gene regulatory states after the initial exposure, which could be consistent with epigenetic regulation. Epigenetic changes involve chromatin modifications that persist after stress and influence gene reactivation, as well as transcriptional bookmarking, where prior stress alters how genes respond to subsequent exposures, sometimes through prolonged association of regulatory factors at stress-responsive promoters. Together, these mechanisms can alter the speed or magnitude of gene re-induction upon subsequent stress exposure (Tehrani *et al*., 2025).

Although DNA methylation, a major epigenetic mechanism in many eukaryotes, is thought to play little or no regulatory role in *Saccharomyces* yeasts, chromatin-based mechanisms of transcriptional memory have been reported in response to repeated stress. In particular, persistent changes in nucleosome organization and promoter architecture have been shown to modify the efficiency with which stress-responsive genes are reactivated after an initial stimulus (D’Urso and Brickner, 2014; D’Urso *et al*., 2016).

Measuring chromatin accessibility (e.g. using ATAC-seq) could help clarify whether memory-like transcriptional responses are mediated by persistent changes in promoter accessibility and therefore explain species-specific patterns of post-desensitization and post-sensitization upon repeated heat wave exposure. Alternatively, repeated heat wave exposure may lead to a more general decline in cellular condition or viability, resulting in transcriptome-wide downregulation of energetically costly processes such as transcription and translation. Such global effects could be particularly pronounced in cold-adapted species and may contribute to the patterns observed for a subset of genes (Fig. 8). While our comparative transcriptomic analyses identify conserved and species-specific pathways associated with repeated heat-wave exposure, future functional studies will be required to determine the causal contribution of these pathways to thermal resilience and transcriptional memory.

## Conclusion

Repeated heat waves elicited a largely conserved core transcriptional response across *Saccharomyces* species, characterized by the induction of proteostasis pathways and the repression of translation and ribosome biogenesis. Yet, despite this shared response, species differed markedly in how this core program was modulated across successive exposures. In particular, memory-like regulatory responses were highly species-specific, with opposite patterns of (de)sensitization in ribosomal and translational pathways observed in species occupying the extremes of thermal tolerance. These contrasting strategies suggest that thermal resilience could be achieved through alternative regulatory routes of a shared transcriptional architecture over time. More broadly, these findings underscore how short-term regulatory dynamics, even in the absence of genetic change, may contribute to performance differences (Zilio *et al*., 2025). Because thermal tolerance shapes fungal survival and host range, warming environments may expand the geographic distribution of thermotolerant species, lower the thermal barrier to mammalian infection, and promote the emergence of novel pathogens (Garcia-Solache and Casadevall, 2010). Together, these results highlight how species-specific regulatory strategies upon exposure to repeated heat waves may influence the ecological resilience and evolutionary potential of species in a changing climate.

## Acknowledgements

Sequencing was supported by a grant from the Royal Physiographic Society of Lund through The Nilsson-Ehle Endowments to CH. The work was funded by the Swedish Research Council (grant 2022-03427) and the Knut and Alice Wallenberg Foundation (grant number 2024.0216) to RS. The authors acknowledge support from the National Genomics Infrastructure in Stockholm funded by Science for Life Laboratory, the Knut and Alice Wallenberg Foundation and the Swedish Research Council, and SNIC/Uppsala Multidisciplinary Center for Advanced Computational Science for assistance with massively parallel sequencing and access to the UPPMAX computational infrastructure.

