## Supplementary Figure S1 for "Repeated heat waves trigger divergent transcriptional responses in cold- and warm-adapted yeast species"

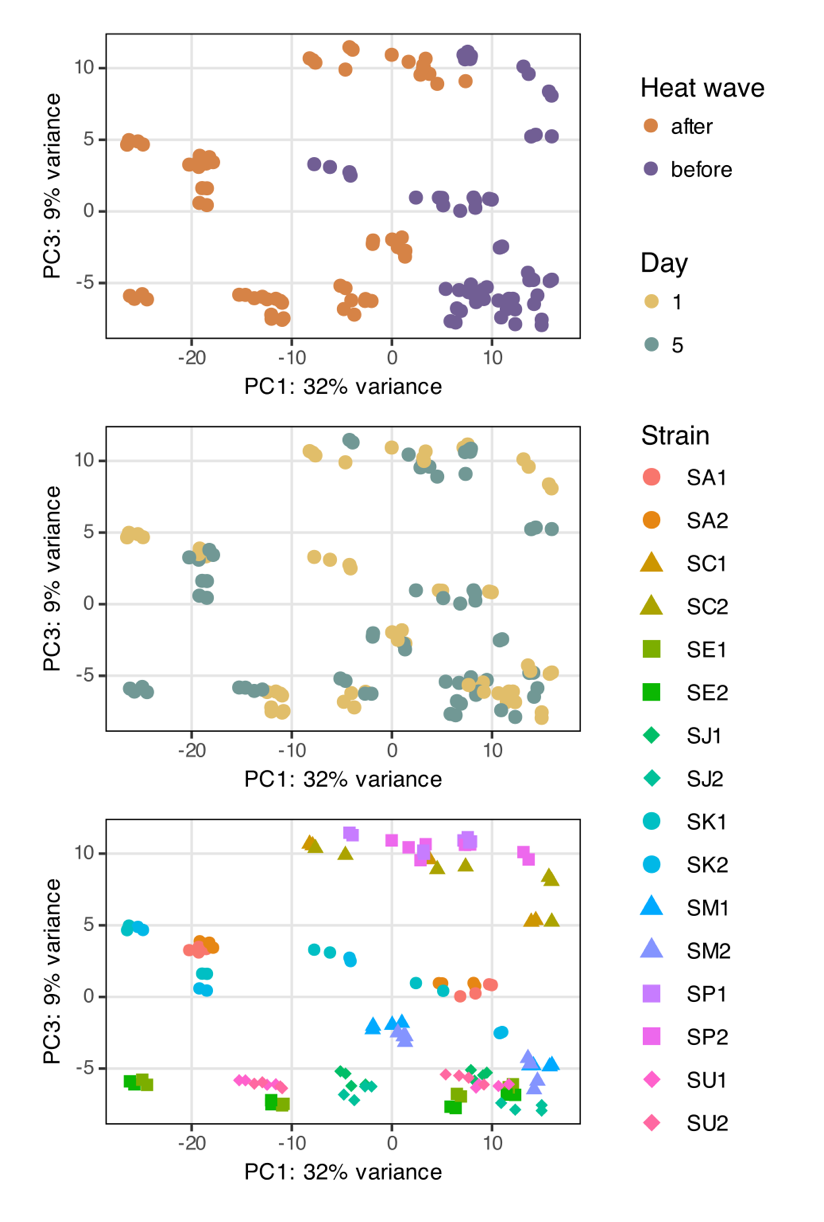

**Supplementary Figure S1.** Projection of all *Saccharomyces* transcriptomes on the first two principal components using PCA. Data points represent the top 500 most variable genes of a sample (default parameter for plotPCA function, out of 2,859 common genes) and are colored *before* or *after* the heat wave (top panel), day 1 or 5 (middle panel), and strain (bottom panel, n = 8 per strain). Read counts for genes were obtained from DESeq2 and processed with *vst*, which computes a variance stabilizing transformation.
